# Slow-Rate Temporal Sampling Deficits During Naturalistic Speech Listening in Children with Developmental Language Disorder

**DOI:** 10.64898/2026.04.16.718920

**Authors:** Mahmoud Keshavarzi, Georgia Feltham, Susan Richards, Lyla Parvez, Usha Goswami

**Affiliations:** Centre for Neuroscience in Education, Department of Psychology, University of Cambridge, Cambridge, CB2 3EB, United Kingdom

## Abstract

Neural tracking of slow temporal modulations in speech supports extraction of prosodic and syllabic structure critical for speech comprehension, yet whether these automatic cortical tracking mechanisms are altered in children with developmental language disorder (DLD) remains unclear. We recorded MEG while children with and without DLD listened to a story, and quantified source-level lagged speech-brain coherence and frequency-specific cortical functional connectivity across bilateral cortical regions. Children with DLD showed significantly reduced coherence in the 0.9–2.5 Hz range associated with prosodic information in the story, spanning bilateral auditory and speech-related cortex. In the 2.5–5 Hz range, linked to syllabic-rate modulations in the story, group differences were right-lateralised. No reliable differences were observed at higher modulation rates (5–9 Hz or 12–40 Hz). These coherence reductions were accompanied by altered functional connectivity between cortical regions across all frequency bands, indicating disrupted large-scale coordination within speech-processing networks in DLD.

## Introduction

Studies in auditory neuroscience show that the brain transforms continuous acoustic speech input into meaningful linguistic structure by using neural mechanisms that operate across multiple temporal scales. Converging evidence from adult studies employing electroencephalography (EEG), electrocorticography (ECoG) and magnetoencephalography (MEG) demonstrates that cortical activity aligns with the temporal dynamics of the speech amplitude envelope (AE), enabling the extraction of prosodic, syllabic and segmental information through hierarchical temporal sampling (multi-time resolution models: Giraud & Poeppel, 2012; Poeppel & Assaneo, 2020).

Adult data suggest that delta-band oscillations (0.5–4 Hz) are preferentially coupled to slow modulations in speech, yielding information about prosody, stress patterns and phrasal boundaries (Giraud & Poeppel, 2012; Gross et al., 2013). Theta-band (4–8 Hz) oscillations synchronise with syllable-rate acoustic fluctuations, facilitating the parsing of continuous speech into syllabic units (Doelling et al., 2014; Ding & Simon, 2014; Luo & Poeppel, 2007). Faster oscillatory activity in gamma band is thought to encode finer-grained phonetic and acoustic detail, supporting rapid analysis within shorter temporal integration windows (Giraud & Poeppel, 2012). This multiscale alignment between neural activity and speech dynamics – often referred to as speech-brain entrainment or cortical tracking – appears critical for speech perception and language processing under naturalistic listening conditions.

Disruptions to these multiscale speech-brain alignment mechanisms may therefore be implicated in developmental language difficulties. Developmental Language Disorder (DLD) is a common neurodevelopmental condition, affecting approximately 7% of children, and is characterised by persistent impairments in language comprehension and production despite adequate hearing, and sufficient language exposure (Leonard, 2017). These linguistic difficulties occur in all languages, and are comprehensive. The CATALISE multi-national study of DLD identified linguistic processing problems with syntax and grammar, a limited understanding of word meanings, impairments in verbal short-term memory, and difficulties with phonology (word sound structure) as core linguistic deficits (Bishop et al., 2017). Given the wide range of linguistic difficulties, the aetiology is likely to involve impairments in fundamental aspects of acoustic processing of the speech signal. An early influential acoustic account proposed that DLD arises, at least in part, from atypical auditory-temporal processing at rapid timescales >40 Hz related to phonemes. Rapid Auditory Processing (RAP) theory (Tallal & Piercy, 1973; Tallal, 1980, 2004) was based on tasks using discrete nonspeech (tone) stimuli rather than continuous speech. Temporal order judgement (TOJ) and other tasks indicated that children with DLD have difficulty processing brief, rapidly successive acoustic events. This was proposed to affect perception of features such as formant transitions and voice onset time that are critical for phoneme identification and categorical speech perception. However, interventions based on RAP such as FastForWord^TM^ have produced mixed results in randomised control trials (Gillam et al., 2008), and have not improved language skills in children with DLD.

By focusing on the phoneme level, RAP may be omitting to consider a more foundational linguistic level regarding language processing. Infants are known to begin the task of acquiring a linguistic system by using speech rhythm rather than by using phonemes (Mehler et al., 1988; Nazzi et al., 1998). Recent neural data suggest infants do not begin to encode phonetic detail until later in the first year of life, and that this encoding develops quite slowly (Di Liberto et al., 2023). Accordingly, investigation of how children with DLD process the low-frequency temporal rates in the speech AE relevant to extracting prosodic structure may be useful. Indeed, recent cortical tracking studies with neurotypical infants aged 4, 7 and 11 months have shown that delta-band cortical tracking is a significant predictor of later language outcomes measured at 2 years (Attaheri et al., 2024). Accordingly, a focus on the accuracy of the alignment of slow cortical oscillations <10 Hz during natural speech processing may offer a complementary framework to RAP for understanding DLD. Such a framework is offered by Temporal Sampling (TS) theory (Goswami, 2011; 2015; 2022). TS theory adopts an acoustic perspective on rhythm based on the speech AE and on the discrimination of amplitude rise times (ARTs). ARTs are rates of transition from lower to higher amplitude that co-occur at multiple rhythmic time frames within speech, which also act as automatic triggers for cortical tracking via phase-resetting oscillatory networks in auditory cortex (Doelling et al., 2014). The perception of ART is impaired in children with DLD (Corriveau et al., 2007; Corriveau & Goswami, 2009; Fraser et al., 2010; Beattie & Manis, 2013; Richards & Goswami, 2015). Accordingly, cortical tracking processes at one or more temporal rates may also show impairments. TS theory proposed that developmental language difficulties arise from atypical oscillatory encoding of slow temporal modulations in the speech amplitude envelope below ∼10 Hz which carry information about speech rhythm, prosodic structure, and syllabic timing (Greenberg, 2012). Therefore, investigation of the processing of slower temporal modulations in speech in children with DLD may be informative.

Although RAP and TS theories are not mutually exclusive and may reflect disruptions at different levels of temporal integration within the auditory system, to date only TS theory has been tested using natural speech (ENGLISH: Attaheri et al., 2022, 2024; Power et al., 2016; Keshavarzi et al., 2022; Mandke et al., 2022; SPANISH: Molinaro et al., 2016; FRENCH: Destoky et al., 2020, 2022). Most TS studies to date are focused on explaining developmental dyslexia, a disorder of written language processing that presents with phonological (speech-sound) difficulties. As noted earlier, phonological difficulties are also a core feature of DLD (Bishop et al., 2017). The prior child dyslexia studies revealed less accurate cortical speech tracking in both the delta (English, Spanish) and theta (French) bands. In the infant cortical tracking studies, individual differences in the accuracy of delta-band tracking were predictive of both vocabulary and phonology (measured by a non-word repetition task). By contrast, infants with more theta power spectral density (PSD) during natural speech listening or with a higher ratio of theta power to delta power had worse language outcomes (Attaheri et al., 2024). These prior developmental data suggest that a focus on both delta-and theta-band neural entrainment during natural speech listening may provide new information about the aetiology of DLD.

Further, a growing body of behavioural and sensory evidence indicates substantial overlap in sensory and linguistic processing difficulties between children with dyslexia and those with DLD (Goswami, 2022). Both developmental disorders are characterised by impairments in the accurate discrimination of ARTs (for a summary of dyslexia data, see Goswami, 2015). ARTs signal rhythmic prominence in speech and both children with DLD and children with dyslexia exhibit atypical perception of syllable stress patterns in words and phrases in experimental tasks (Richards & Goswami, 2015; Cumming et al., 2015a, 2015b). Recently, it has been shown that both groups of children are also characterised by the atypical production of syllable stress patterns when copying aloud multisyllabic words (Keshavarzi et al., 2024a; Parvez et al., 2024). These convergent sensory and behavioural findings indicate possible overlapping origins in low-frequency temporal sampling mechanisms, which are investigated here in DLD.

Surprisingly, direct neurophysiological evidence for atypical cortical tracking of continuous naturalistic speech by children with DLD remains limited. A recent MEG study using spoken word stimuli reported altered temporal dynamics of speech tracking in children with DLD, characterised by reduced maintenance of speech envelope representations at longer latencies relative to typically developing peers (Nora et al., 2024). Importantly, this study focused on isolated words rather than continuous speech, and interpreted the observed effects in terms of impaired retention of acoustic-phonetic information rather than deficits in early stimulus-locked encoding. The majority of neural studies of children with DLD employ fMRI, and report reduced activation in left frontal and temporal cortical areas in the DLD group, but do not throw light on underlying mechanisms (Asaridou & Watkins, 2022, for recent review). Prior MEG studies using speech processing tasks are consistent with the fMRI literature, identifying atypical left hemisphere function particularly in temporal regions (Helenius et al., 2009, 2014). Some fMRI studies also report atypical right hemisphere activation, although a recent review concluded that laterality differences do not seem as fundamental as once thought (Parker et al., 2022). By contrast, prior electrophysiological studies have largely relied on highly controlled or artificial stimuli, or have focused on scalp-level measures, making it difficult to determine whether speech processing is altered at the level of distributed cortical sources within auditory and speech-language networks. Consequently, it remains unclear whether neural abnormalities in DLD primarily affect specific temporal scales of speech processing, such as those associated with prosodic and syllabic structure, or whether they reflect more widespread disruptions in temporal alignment and coordination across the cortical hierarchy during continuous speech perception.

Accordingly, motivated by TS theory, we here recorded MEG in children with and without DLD during a story listening task and focused our analyses on a set of *a priori* defined bilateral auditory and speech-language regions of interest (ROIs). Our selection of ROIs was based on a recent sEEG (stereoelectroenchephalography) study of cortical processing of prosody by adults (Gnanateja et al., 2025) and the adult MRI data regarding prosody reported by Sammler et al. (2015). The ROIs included Heschl’s gyrus (HG), superior temporal gyrus (STG), middle temporal gyrus (MTG), superior temporal sulcus (STS), planum temporale (PT), inferior frontal gyrus pars opercularis (IFGop) and pars triangularis (IFGtri), as well as supramarginal gyrus (SMG), angular gyrus (AG) and inferior parietal lobule (IPL). To examine cortical tracking at different linguistic scales, we used a children’s story for which previously-applied computational modelling had identified the temporal bandings corresponding to prosodic, syllabic and phonemic information respectively (Mandke et al., 2022). In a study of children with dyslexia, Mandke et al. (2022) had analysed neural tracking within four functionally meaningful amplitude-modulation (AM) ranges in the speech envelope of the story. AM rates spanning ∼0.9–2.5 Hz corresponded to prosodic and phrasal-level structure, reflecting the timing of intonational contours and phrasal grouping; AM rates spanning ∼2.5–5 Hz aligned with syllabic timing; a faster modulation rate (spanning ∼5–9 Hz) indexed faster segmental and sub-syllabic structure; and broadband activity in the 12–40 Hz range captured cortical dynamics related to phonemic information. This stimulus-specific and band-specific framework enabled us to test a core tenet of TS theory, namely that atypical language development in DLD is associated with selective disruption of slow temporal sampling mechanisms, rather than a uniform deficit across all timescales of speech processing. The participating children listened to an extended spoken narrative (10 minutes) under natural listening conditions while MEG was recorded (Fig. 1a). MEG data were reconstructed at the cortical level using individual structural MRI (Fig. 1b). Source-level lagged speech-brain coherence was quantified between cortical activity and the speech amplitude envelope across multiple temporal modulation ranges, as well as frequency-specific functional connectivity between cortical regions (Fig. 1c). By identifying multiple ROIs *a priori*, this approach enabled us to assess both local cortical tracking of speech rhythms and the large-scale coordination of distributed speech-processing networks across temporal scales.

**Fig. 1:**
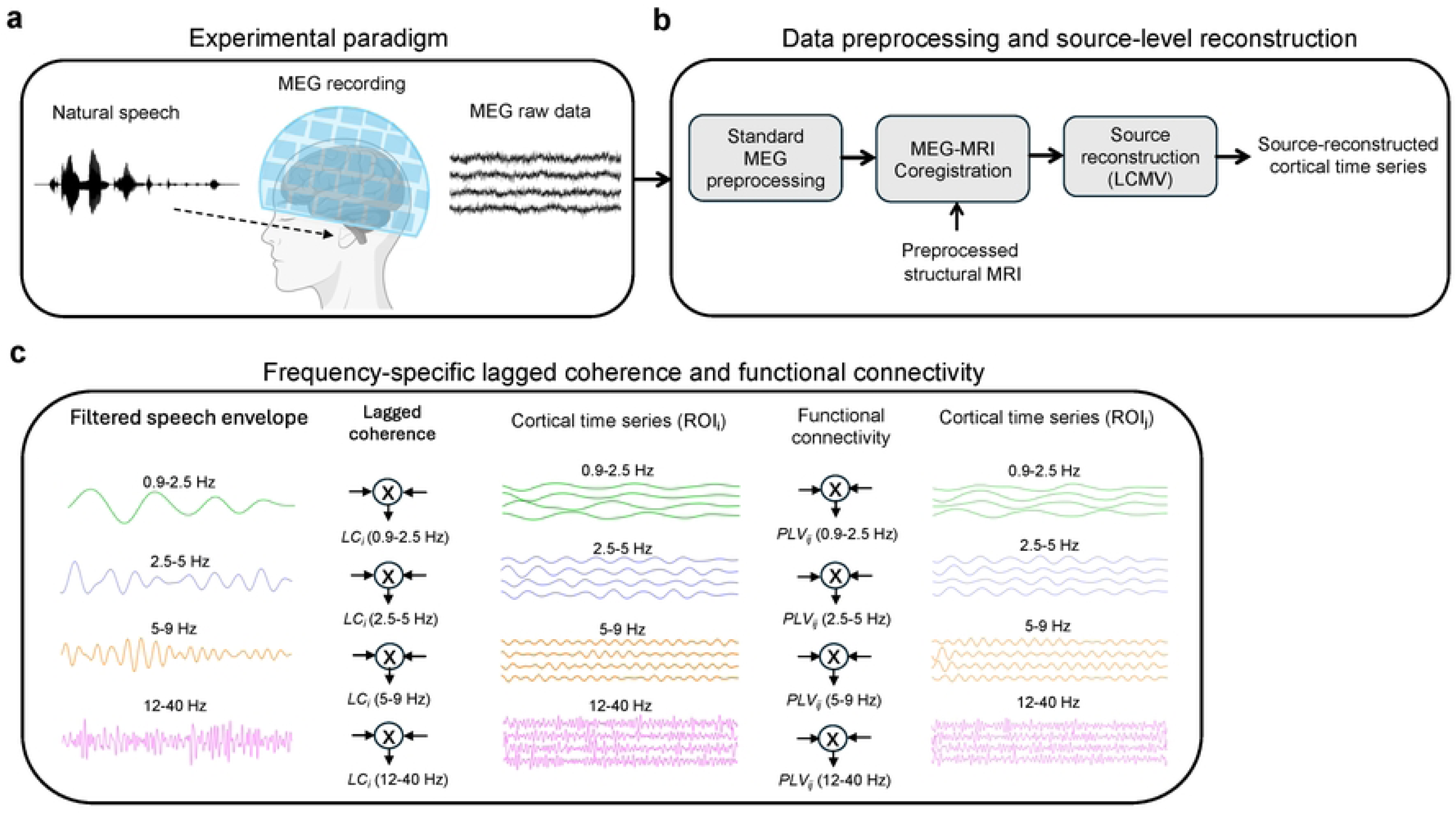
Experimental paradigm, standard preprocessing steps and analysis pipeline for frequency-specific speech-brain lagged coherence and functional connectivity. (a) Participants listened to continuous natural speech while MEG was recorded. (b) MEG data were preprocessed using standard procedures and coregistered with individual structural MRI scans. Source-level cortical time series were reconstructed using an LCMV beamformer. (c) Frequency-specific analyses of speech-brain coherence and cortical functional connectivity. The speech envelope and source-level cortical time series were filtered into four non-canonical frequency ranges (0.9–2.5, 2.5–5, 5–9 and 12–40 Hz). Lagged coherence (*LC_i_*) was computed between the band-limited speech envelope and the time series of each cortical region of interest (ROI_i_). Functional connectivity was then estimated between pairs of cortical regions using phase-locking value (*PLV_ij_*) within the same frequency ranges. Indices *i* and *j* denote cortical ROIs.

We hypothesised that children with DLD would show selective impairments in low-frequency neural tracking of speech, particularly at modulation rates associated with prosodic and phrasal-level structure (0.9–2.5 Hz). We also predicted selective impairments at syllabic rates (2.5–5 Hz). However, no differences at faster temporal scales were expected. Based on neurophysiological TS-driven evidence implicating atypical right-hemisphere processing of connected speech in dyslexia (Di Liberto et al., 2018), we further predicted that group differences would be most pronounced within right-hemisphere temporal and temporo-frontal ROIs, including HG, STG, MTG, STS and PT, as well as their functional interactions with right IFGop and IFGtri. Finally, we hypothesised that reduced speech-brain coherence at slow temporal rates would be accompanied by altered low-frequency functional connectivity between these ROIs, reflecting disrupted large-scale coordination within cortical speech-language networks during naturalistic speech perception in DLD.

## Results

We quantified neural tracking of different temporal components of the speech signal by computing source-level lagged coherence between MEG activity and the band-limited speech envelope (0.9–2.5 Hz, 2.5–5 Hz, 5–9 Hz and 12–40 Hz). For each child, continuous MEG story recordings were reconstructed at the cortical surface using LCMV beamforming, and lagged coherence was estimated using Hilbert-based analytic signals. Coherence values were subsequently averaged within 20 bilateral anatomical ROIs for statistical analysis. Fig. 2a–d shows group-averaged lagged coherence values for each frequency band, plotted separately for left-and right-hemisphere ROIs, together with permutation-derived null-model baselines. Between-group differences were assessed using Wilcoxon rank-sum tests, with statistical interpretation restricted to effects that additionally exceeded the permutation-derived null distributions.

**Fig. 2:**
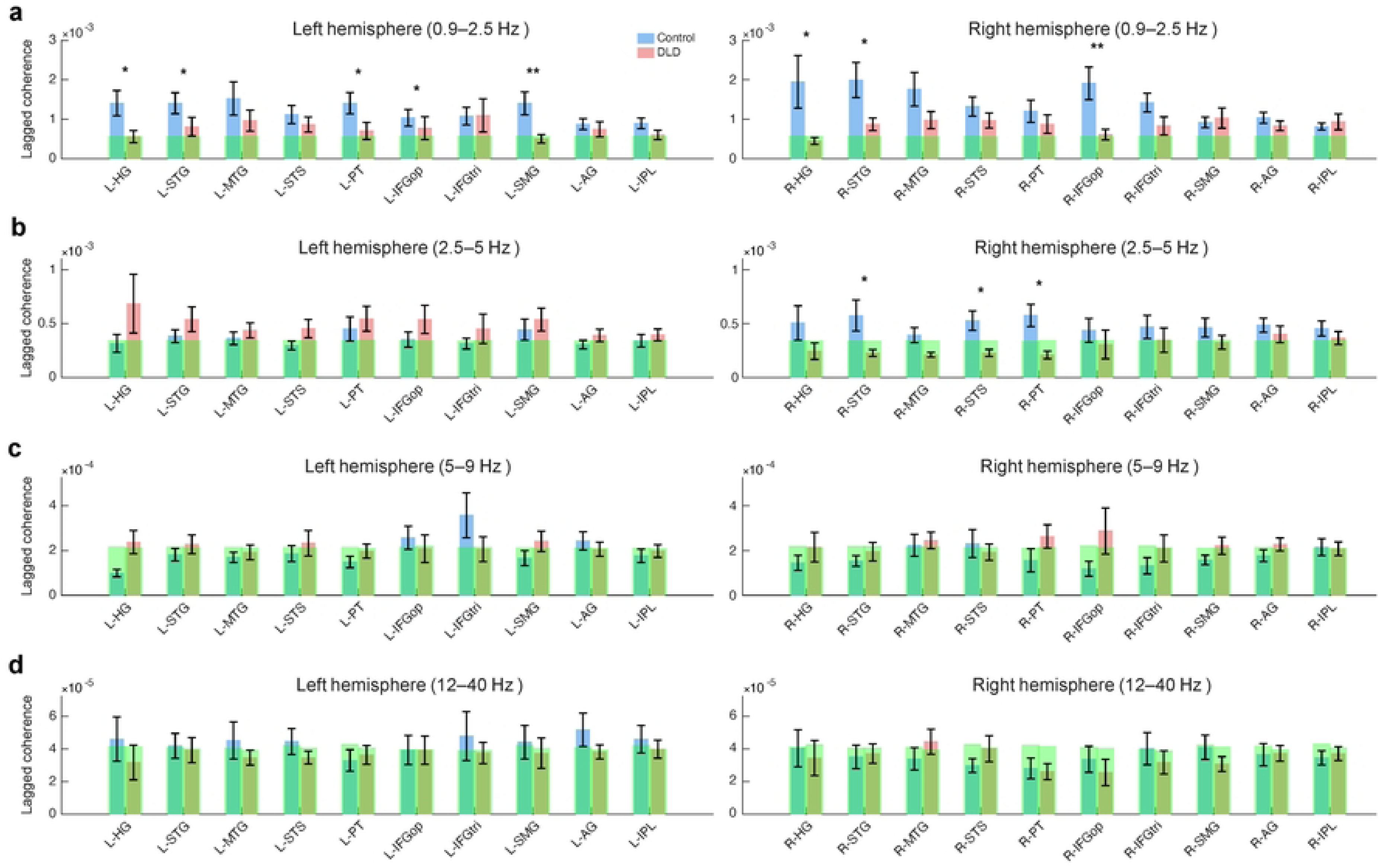
Lagged speech-brain coherence across temporal modulation bands and cortical ROIs. (a–d) Mean lagged coherence between MEG source activity and the band-limited speech envelope is shown for Control (light blue) and DLD (light red) groups across left and right hemisphere ROIs. Each row corresponds to a temporal modulation band: (a) 0.9–2.5 Hz, (b) 2.5–5 Hz, (c) 5–9 Hz and (d) 12–40 Hz. Error bars indicate the standard error of the mean. Narrow green bars behind each group bar represent the mean of the permutation-derived null distribution for the corresponding ROI and frequency band, providing a descriptive baseline estimate of chance-level coherence. Asterisks denote ROIs showing significant group differences based on Wilcoxon rank-sum tests (* *p* < 0.05, ** *p* < 0.01); Wilcoxon results were interpreted only when effects additionally exceeded the permutation-derived null model. Group differences were observed in the 0.9–2.5 Hz and 2.5–5 Hz bands with no reliable differences at higher modulation frequencies.

### Reduced neural tracking of speech prosodic cues in children with DLD

We first examined neural tracking in the 0.9–2.5 Hz band, which reflected slow temporal modulations in our story supporting prosodic and phrasal-level structure. As shown in Fig. 2a, children with DLD exhibited significantly reduced coherence across a distributed bilateral network. Reduced tracking was observed in L-HG (*z* = 2.41, *p* = 0.016), L-STG (*z* = 2.14, *p* = 0.033), L-PT (*z* = 2.41, *p* = 0.016), L-IFGop (*z* = 2.04, *p* = 0.041) and L-SMG (*z* = 3.33, *p* < 0.001). Significant reductions were also present in the right hemisphere, including R-HG (*z* = 2.14, *p* = 0.033), R-STG (*z* = 2.27, *p* = 0.023) and R-IFGop (*z* = 3.23, *p* = 0.001). In all cases, coherence values exceeded the permutation-derived null model, demonstrating reliable above-chance speech-brain coupling and robust group-level differences. These findings indicate a widespread deficit in slow-timescale neural tracking in DLD across bilateral auditory and speech-related cortices (see Fig. 3a–b).

**Fig. 3:**
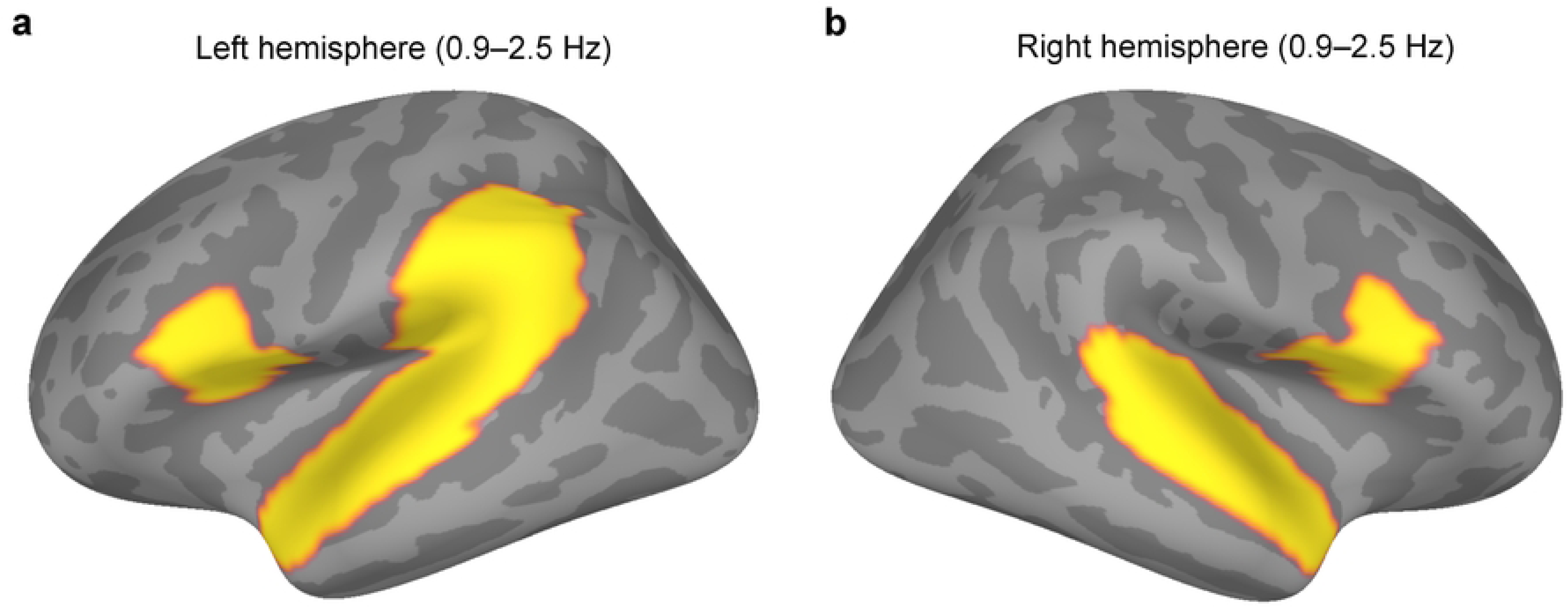
Reduced prosodic-rate (0.9–2.5 Hz) speech-brain coherence in developmental language disorder. (a) Left hemisphere; (b) Right hemisphere. Cortical surface maps show regions exhibiting reduced speech-brain lagged coherence in children with DLD relative to typically developing controls in the 0.9–2.5 Hz band. Group differences were observed across a distributed bilateral network encompassing auditory and speech-related regions, including Heschl’s gyrus (HG), superior temporal gyrus (STG), planum temporale (PT) and inferior frontal gyrus pars opercularis (IFGop), as well as the supramarginal gyrus (SMG). Only regions in which group differences additionally exceeded the permutation-derived null model are shown. Maps are displayed on the fsaverage inflated cortical surface (lateral views) and are thresholded for visualisation.

### Syllabic-rate tracking differences are restricted to right temporal cortex

In the 2.5–5 Hz band (Fig. 2b), corresponding to syllabic-rate temporal structure in our story, Wilcoxon rank-sum tests indicated reduced lagged coherence in children with DLD in several right-hemisphere ROIs, including R-STG (*z* = 2.37, *p* = 0.018), R-MTG (*z* = 2.23, *p* = 0.026), R-STS (*z* = 2.41, *p* = 0.016) and R-PT (*z* = 2.55, *p* = 0.011).

However, when benchmarked against the permutation-derived null model, only the reductions in R-STG, R-STS and R-PT exceeded chance-level coherence. The effect in R-MTG did not exceed the null distribution and was therefore not interpreted further. No significant effects were observed in left-hemisphere ROIs. Together, these findings indicate a right-lateralised but spatially restricted alteration of syllabic-rate speech tracking in DLD (see Fig. 4a–b).

**Fig. 4:**
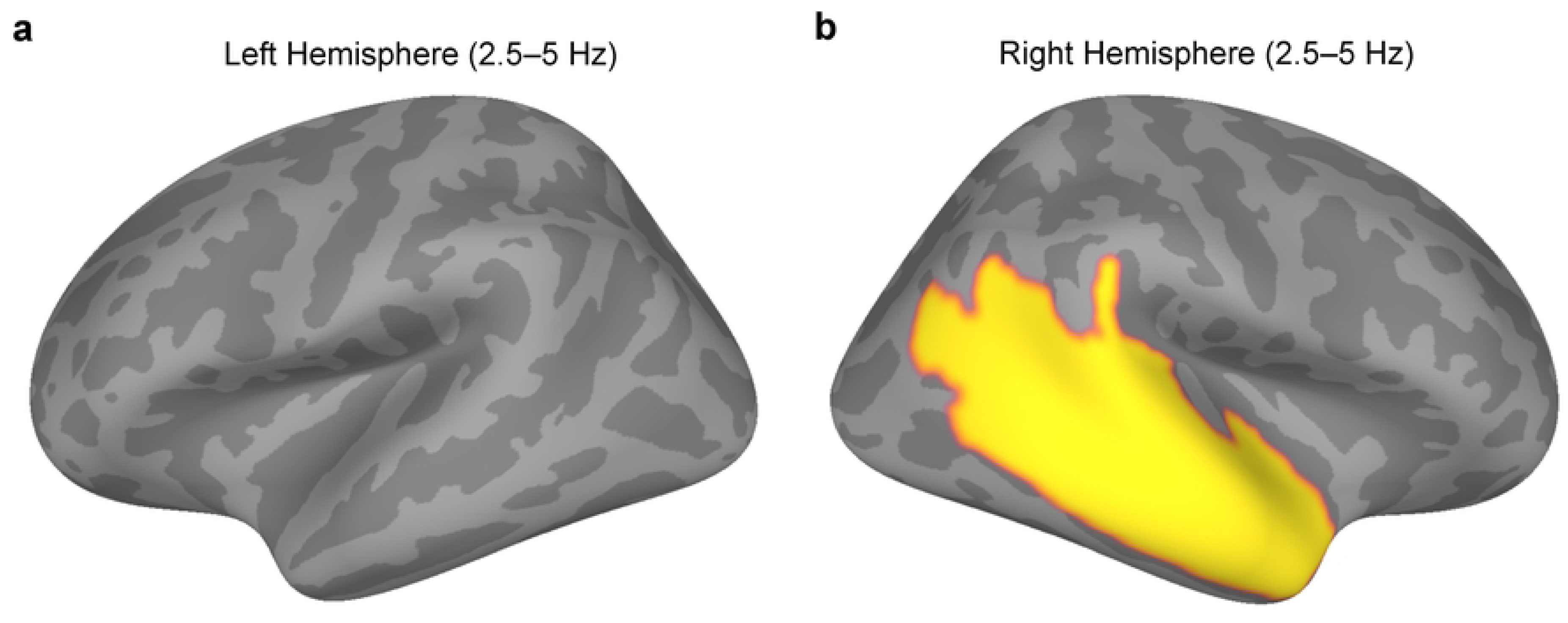
Right-lateralised reductions in syllabic-rate (2.5–5 Hz) speech-brain coherence in developmental language disorder. (a) Left hemisphere; (b) Right hemisphere. Cortical surface maps show regions exhibiting reduced speech-brain lagged coherence in children with DLD relative to typically developing controls in the 2.5–5 Hz band. Group differences were spatially restricted and confined to right-hemisphere temporal regions, including the superior temporal gyrus (STG), superior temporal sulcus (STS) and planum temporale (PT). Regions exhibiting nominal group differences that did not exceed the permutation-derived null model are shown in grey and were not interpreted further. Maps are displayed on the fsaverage inflated cortical surface (lateral views) and are thresholded for visualisation.

### No reliable group differences at higher modulation frequencies

Finally, we examined the 5–9 Hz and 12–40 Hz bands, which corresponded to segmental and higher-frequency AMs in our story. Although a Wilcoxon rank-sum test identified a group difference in R-PT (*z* = –2.14, *p* = 0.033) in the 5–9 Hz band, this effect did not exceed the permutation-derived null model and was therefore not interpreted further. As shown in Fig. 2c–d, no ROIs showed reliable group differences after benchmarking against the permutation-derived null distributions. These results indicate that neural tracking impairments in children with DLD are selective to slower temporal rates, particularly those that encode prosodic and phrasal-level structure, with weaker and more variable differences at syllabic timescales and no evidence for systematic disruptions at faster temporal modulations.

### Altered functional connectivity across frequency bands in children with DLD

We next examined frequency-specific functional connectivity during continuous speech listening by computing the phase-locking value (PLV) between pairs of regional time series (Fig. 5a–d). In contrast to the speech–brain coherence analyses, group differences in functional connectivity were evident across all frequency bands, indicating alterations in inter-regional phase synchronisation in children with DLD that extended beyond low-frequency stimulus tracking.

**Fig. 5:**
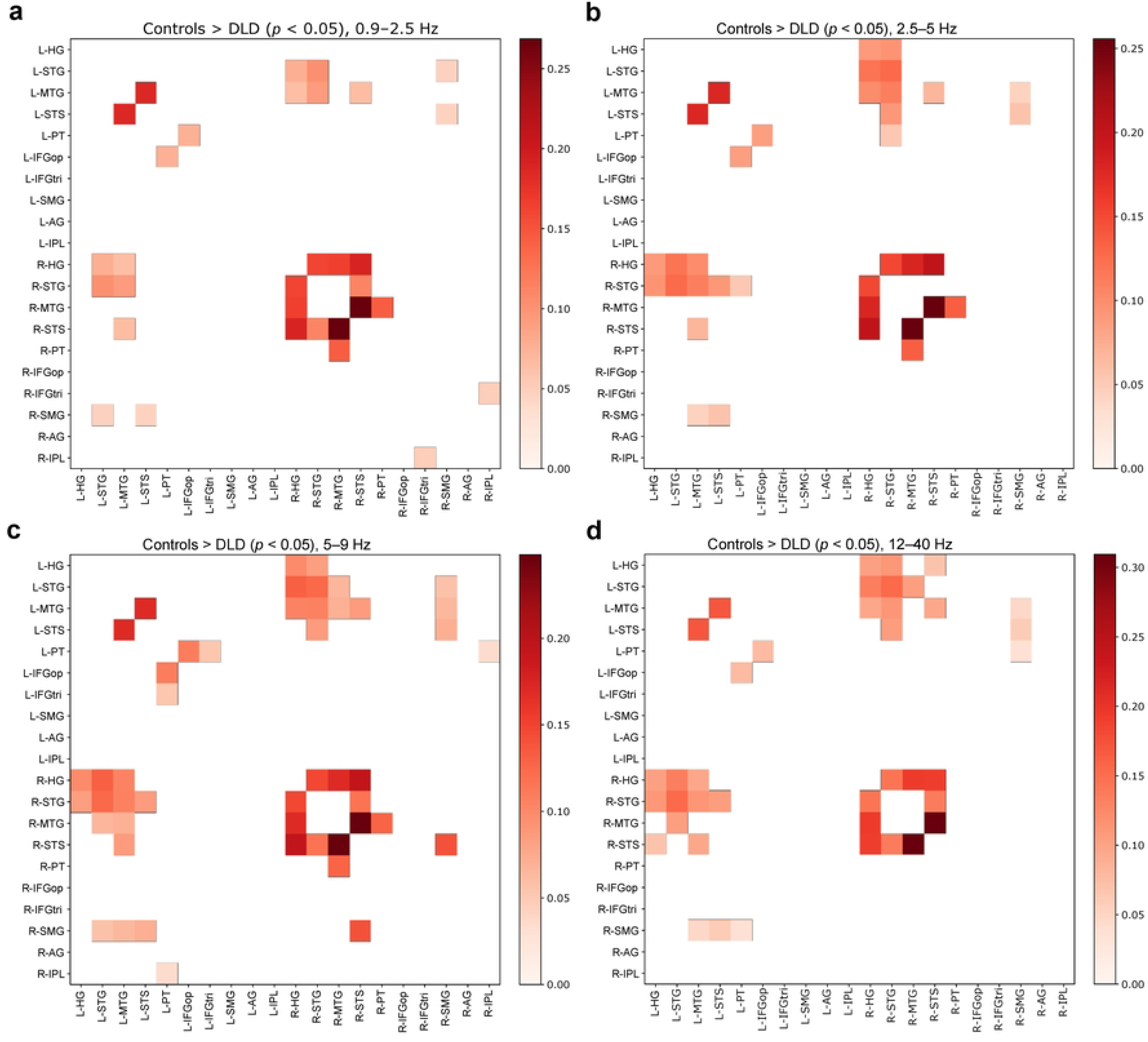
Group differences in frequency-specific functional connectivity during natural speech listening. (a–d) Matrices show significant differences in phase-locking value (PLV) between typically developing control children and children with DLD (Controls − DLD) across cortical ROIs for four frequency ranges: (a) 0.9–2.5 Hz, (b) 2.5–5 Hz, (c) 5–9 Hz and (d) 12–40 Hz. Only ROI pairs exhibiting significant group differences (*p* < 0.05, permutation-based testing) are displayed; non-significant connections are masked. Warmer colours indicate stronger functional connectivity in controls relative to DLD. ROIs are ordered identically along both axes, with left-and right-hemisphere regions indicated by L-and R-prefixes, respectively.

In the 0.9–2.5 Hz band, significant group differences were observed across 16 ROI pairs, forming a distributed network spanning bilateral temporal and temporo-frontal regions. The most connected node was R-HG, which participated in five altered connections, indicating hub-like involvement at prosodic timescales. Additional prominent nodes included R-STG, R-STS, and L-MTG (four connections each), with further contributions from bilateral superior temporal and parietal regions. This pattern reflects a bilateral and integrative network architecture, with a right-temporal emphasis but substantial engagement of left temporal cortex, consistent with a large-scale functional difference when encoding prosodic and phrasal-level speech structure. This functional difference appears to originate in primary auditory cortex.

In the 2.5–5 Hz band, group differences involved 18 ROI pairs and exhibited a more focal and hierarchical network organisation. The highest degree of involvement was observed in R-HG and R-STG, each participating in six altered connections, indicating hub-like involvement of right auditory and superior temporal cortex at syllabic rates. In addition, L-MTG emerged as a prominent secondary node with five connections, suggesting bilateral engagement of middle temporal regions. Other areas showed substantially fewer connections, resulting in a tiered network structure rather than the distributed pattern observed at the slower temporal rate (0.9–2.5 Hz). In the 5–9 Hz band, group differences were observed across 24 ROI pairs, representing the most extensive connectivity reorganisation among all frequency ranges examined. The network was characterised by multiple hub-like nodes, including R-HG, R-STG, and L-MTG (six connections each), with additional prominent involvement of R-MTG and R-STS (five connections each). Compared with lower-frequency bands, altered connectivity at 5–9 Hz engaged a broader set of temporal regions and showed a pronounced right-hemisphere bias, while retaining strong participation of left middle temporal cortex. Notably, these widespread connectivity changes occurred despite the absence of corresponding speech-brain coherence differences at this frequency, suggesting that the altered inter-regional synchronisation reflects changes in large-scale network dynamics rather than impaired tracking of specific speech rhythms.

In the 12–40 Hz band, significant group differences were identified for 20 ROI pairs, indicating a structured but reduced network reorganisation relative to the 5–9 Hz band. The highest degree of altered connectivity again involved R-HG and R-STG (six connections each), with secondary hub-like contributions from L-MTG and R-STS (five connections each). Several additional temporal regions exhibited moderate connectivity, yielding a compact and organised network centred on auditory and superior temporal cortex. As with the 5–9 Hz band, these effects were observed in the absence of speech-brain coherence differences, indicating that higher-frequency connectivity alterations in DLD are not driven by impaired speech tracking but instead reflect persistent differences in inter-regional coordination during receptive speech processing.

## Discussion

The present study provides source-level neurophysiological evidence that children with DLD exhibit selective impairments in the cortical tracking of slow temporal structure (<5 Hz) in naturalistic speech. No impairments were detected in cortical tracking at higher frequencies above 5 Hz. Reduced cortical tracking at lower frequencies could be expected to affect the extraction of both prosodic and syllable-level information in the children’s story used as the stimulus. Reduced tracking was accompanied by frequency-dependent alterations in large-scale cortical functional connectivity which spanned both hemispheres and all frequency bands. These coherence deficits coupled with widespread atypicalities in inter-regional functional connectivity indicate that atypical speech processing in DLD reflects not only local impairments in speech-brain coupling but also altered coordination within distributed cortical speech networks.

The most important finding of the present study is the reduction in speech-brain coherence in the 0.9–2.5 Hz range across bilateral auditory and speech-related cortex in children with DLD. This frequency range corresponds to the slow amplitude modulations that carry information about prosodic phrasing, intonational contours, speech rhythm and hierarchical grouping in speech (Greenberg, 2012; Ghitza & Greenberg, 2009; Giraud & Poeppel, 2012; Gross et al., 2013). This is also the frequency range most affected in children with developmental dyslexia (Goswami, 2022, for review), as well as the frequency range showing the highest levels of tracking accuracy in infants aged <12 months (Attaheri et al., 2022, 2024). At least for a stress-timed language like English, therefore, the efficiency of cortical tracking at ∼2 Hz appears to be foundational in developing a well-functioning linguistic system. Reduced coherence in the 0.9–2.5 Hz range was observed in bilateral HG, STG, PT and IFGop, indicating that atypical processing at this timescale is not confined to early auditory cortex but extends across distributed speech-processing networks. These extensive frequency-specific differences in functional connectivity may help to explain why children with DLD experience difficulties across all the linguistic domains of syntax, phonology and semantics (Bishop et al., 2017). In addition to these prosodic-rate effects, group differences were also observed at the rates denoting syllabic information in the story (2.5–5 Hz). Interestingly, these differences were more spatially restricted and were largely confined to right-hemisphere temporal regions. In the 2.5–5 Hz band, reduced coherence in children with DLD was evident primarily in R-STG, R-STS and R-PT, consistent with some of the prior fMRI data indicating right-hemisphere neural processing atypicalities (Parker et al., 2022).

Together, these findings indicate a graded impairment in low-frequency speech-brain alignment in DLD, with the strongest and most spatially extensive effects found at prosodic timescales and weaker, right-lateralised effects found at syllabic rates. These findings provide direct neurophysiological support for the TS framework, which proposes that atypical language development arises from impaired cortical encoding of the slow temporal modulations in speech that support both prosodic and syllabic structure (Goswami, 2011, 2015, 2022). The graded nature of these effects – widespread and bilateral at prosodic rates but right-lateralised at syllabic rates – suggests that TS impairments in DLD are not limited to a single temporal scale but span multiple levels of the speech rhythm hierarchy. The right-hemisphere predominance of the syllabic-rate effects observed in the 2.5–5 Hz band are also consistent with the hemispheric asymmetries in temporal integration proposed by the Asymmetric Sampling in Time (AST) hypothesis (Poeppel, 2003; Boemio et al., 2005), by which slower-rate temporal information is preferentially processed by the right hemisphere. However, AST did not consider rates slower than the syllabic rate.

No reliable group differences were observed in speech-brain coherence at higher modulation frequencies (5–9 Hz and 12–40 Hz), which are thought to correspond to faster segmental and sub-segmental temporal structure (Giraud & Poeppel, 2012). On a developmental analysis, this may not be surprising, as behavioural studies suggest that children’s knowledge about segmental units develops relatively slowly and is related to learning to read (Ziegler & Goswami, 2005). In the current study, faster cortical tracking rates did not differentiate children with DLD from control children. Regarding RAP theory, these findings suggest that neural tracking of gamma-band acoustic fluctuations (>40 Hz, the frequency rate foregrounded by RAP theory as relevant for phonemic knowledge) is not impaired in children with DLD. Neural tracking of more rapid speech information appears largely preserved during naturalistic speech listening, at least on the basis of the methods used in our study. This result requires replication, but may indicate that the origins of DLD lie not with the phoneme but rather with the syllabic rhythm patterns that comprise prosody.

In contrast to the speech-brain coherence results, functional connectivity analyses revealed group differences across all frequency bands. This finding is consistent with the elevated EEG band-power that appears to characterise the DLD brain when listening to repetition of the syllable “ba” (Keshavarzi et al., 2024b), and implies widespread alterations in inter-regional phase synchronisation. With respect to the slowest temporal rate (0.9–2.5 Hz), children with DLD showed reduced functional connectivity across a distributed bilateral network encompassing auditory and temporal regions, with additional involvement of temporo-parietal and inferior frontal cortex. Connectivity differences were evident in both hemispheres and included a mixture of intra-hemispheric and inter-hemispheric connections, consistent with disrupted large-scale coordination at prosodic timescales. Within this network, right auditory and superior temporal regions, particularly R-HG, R-STG and R-STS, showed prominent involvement, while L-MTG also contributed substantially, indicating that impaired prosodic-rate processing in DLD engages a bilateral but right-weighted temporal network rather than exhibiting a focal locus of dysfunction. Given the established role of low-frequency phase synchronisation in supporting long-range cortical integration (Varela et al., 2001), this pattern suggests that the coordinated integration of slow rhythmic and phrasal-level speech information across distributed cortical systems is inefficient in the DLD brain.

At syllabic rates (2.5–5 Hz), functional connectivity differences were more focal and hierarchically organised than at the slowest timescale. Altered connectivity was dominated by right auditory and superior temporal cortex, with R-HG and R-STG emerging as the most connected nodes and L-MTG acting as a prominent secondary contributor. Compared with the distributed pattern observed at 0.9–2.5 Hz, this tiered organisation suggests a sharpening of network disruption at syllabic timescales, centred on regions critical for processing speech rhythm and timing. The close correspondence between these network-level alterations and the right-lateralised coherence differences observed at 2.5–5 Hz supports the interpretation that atypical syllabic-rate processing in DLD reflects both reduced local neural alignment and weakened inter-regional coordination within temporally specialised speech networks. Importantly, group differences in functional connectivity persisted at higher frequencies (5–9 Hz and 12–40 Hz) despite the absence of corresponding group differences in speech–brain coherence. At these frequencies, connectivity alterations were extensive at 5–9 Hz and more constrained but still structured at 12–40 Hz, with repeated involvement of right auditory and superior temporal regions alongside bilateral temporal cortex. This dissociation indicates that preserved local tracking of faster speech modulations does not necessarily entail efficient large-scale coordination across cortical networks. One implication is that, while neural encoding of segmental and high-frequency acoustic information may be largely intact in DLD, the integration of these representations across distributed cortical systems remains atypical. Such a decoupling between local stimulus tracking and inter-regional synchronisation accords with frameworks emphasising communication through coherence (Varela et al., 2001; Fries, 2005, 2015), in which phase alignment across cortical areas regulates the efficiency and stability of information transfer during complex cognitive tasks, including naturalistic speech processing. The atypical network-level organisation demonstrated here may contribute to variability in language outcomes and help to explain why difficulties in speech comprehension and language use persist in DLD despite relatively intact performance on some basic auditory psychophysical measures with tones (e.g., backward masking and frequency-modulation detection) in quiet listening conditions (Bishop et al., 1999).

In summary, here we show that DLD is associated with selective impairments in slow-timescale cortical speech tracking and frequency-dependent disruptions of cortical functional connectivity during continuous speech listening. By linking reduced prosodic-and syllabic-rate neural alignment to altered large-scale network coordination – particularly within right-hemisphere temporal and temporo-frontal systems – the findings provide converging mechanistic support for some prior fMRI data (Helenius et al., 2014, 2009; Asaridou & Watkins, 2022). More broadly, the results highlight slow temporal sampling as a feature of DLD, supporting TS theory, and identifying novel neurophysiological targets for future interventions.

## Materials and Methods

### Participants

Twenty-eight children took part in the study, including 14 typically developing control children (mean age = 9.31 years, S.D. = 0.47) and 14 children with DLD (mean age = 9.60 years, S.D. = 1.02). All participants were native English speakers with no history of neurological disorders. All participants were taking part in an ongoing study of auditory processing in DLD, and those children with suspected language difficulties were nominated by the special educational needs teachers in their schools. Children in the TD group were nominated by classroom teachers as being typically-developing. All participants exhibited normal hearing when tested with an audiometer. In a short hearing test across the frequency range 0.25 – 8 kHz (0.25, 0.5, 1, 2, 4, 8 kHz), all children were sensitive to sounds within the 20 dB HL range. Language status was ascertained by a trained speech and language therapist, who administered two subtests of CELF-V (Wiig et al., 2017) to all the participants: recalling sentences and formulating sentences. Those children who appeared to have language difficulties then received (depending on age) two further CELF subtests drawn from word structure, sentence comprehension, word classes and semantic relationships. Children who scored at least 1 S.D. below the mean (7 or less when the mean score is 10) on at least 2 of these 4 subtests were included in the DLD group. MRI scans were available for all individuals except five participants, for whom the FreeSurfer fsaverage template was used during source modelling. All participants and their parents provided informed consent to take part in the study, in accordance with the Declaration of Helsinki. The study was reviewed by the University of Cambridge, Psychology Research Ethics Committee and received a favourable opinion.

### Stimuli and Experimental Design

Participants listened to a continuous naturalistic narrative (“Iron Man”), divided into 10 sequential ∼1-min segments. MEG data were collected during two separate recording sections (each lasting approximately 5 minutes), with five story segments presented in each section. Each story segment was followed by a 2-s silent pause. Speech stimuli were sampled at 44.1 kHz. Onset times for each story part were logged and later aligned precisely to MEG time series for coherence analysis.

### MEG Data Acquisition

MEG data were acquired using a 306-channel Elekta Neuromag system (102 magnetometers and 204 planar gradiometers) in a magnetically shielded room. Signals were sampled at 1000 Hz with an online band-pass filter (0.1–330 Hz). Head position indicator (HPI) coils were used to track head position, and standard fiducials (nasion and left/right preauricular points) and scalp points were digitised for co-registration. Two-minute empty-room recordings were acquired for noise covariance estimation using the same acquisition parameters as the task recordings. To ensure attention to the narrative, children were informed prior to testing that they would answer simple comprehension questions following each story section. Comprehension was assessed using four questions in total (score range: 0–4). Mean scores (± S.D.) were 3.64 ± 0.50 for the TD group and 3.57 ± 0.65 for the DLD group. A Wilcoxon rank-sum test revealed no significant group difference (*z* = 0.11, *p* = 0.91), indicating comparable comprehension performance across groups.

### MEG data preprocessing

MEG preprocessing was performed in Python using MNE-Python (Gramfort et al., 2013). For each child, environmental and movement-related noise were suppressed using Maxwell filtering (signal space separation). The data were then band-pass filtered between 0.2 and 48 Hz with zero-phase FIR filtering and resampled to 200 Hz, using MNE-Python (Gramfort et al., 2013). Power spectral density estimates and time domain of MEG channel were inspected to identify excessively noisy channels. Each participant had a manually curated list of bad sensors, which were marked in the data structure and subsequently interpolated using MNE’s spherical spline interpolation. Residual physiological and movement artifacts were removed using independent component analysis (ICA) implemented in MNE-Python (Gramfort et al., 2013). A copy of the data was additionally high-pass filtered at 1 Hz and used to fit an Infomax ICA decomposition. For each child, artefactual components (eye blinks, cardiac activity, movement and sensor noise) were identified manually based on their spatial topographies, time courses and power spectra, and were excluded using participant-specific IC index lists. The cleaned ICA solution was then applied to the 0.2–48 Hz filtered continuous MEG data. All preprocessed recordings were visually inspected to ensure artifact removal and signal integrity, and the resulting cleaned datasets were saved for subsequent source reconstruction and coherence analyses. All recordings, including the empty-room recording, were resampled to 200 Hz.

### Construction of continuous story MEG time series

Each child contributed two preprocessed MEG runs containing five story segments interleaved with 2 s silent intervals. To create a continuous “story” time series, we used the recorded onset times and measured durations to crop each segment from the raw runs. For each run, five segments were extracted by cropping from the onset of each story part to onset + duration, then advancing the start time by the segment duration plus 2 s to skip the silent interval. Segments from the first and second runs were concatenated into a single continuous recording. To ensure consistent geometry, the device-to-head transform of the second run was set equal to that of the first before concatenation. The concatenated story recording was then used for analysis, ensuring a matched duration across children and alignment with the speech envelope.

### Stimulus envelope calculation

Speech stimuli consisted of ten ∼1-minute segments from a narrated “Iron Man” story. For analysis, speech stimuli were converted to 64-bit floating-point format and concatenated to form a single continuous waveform sampled at 44.1 kHz. We first computed a broadband amplitude envelope by taking the magnitude of the analytic signal obtained via the Hilbert transform. The resulting envelope was normalised by its maximum value. To align the speech envelope with the MEG data and reduce computational load, the envelope was then resampled to 200 Hz using Fourier-based resampling, which includes appropriate anti-alias filtering. This envelope was subsequently band-pass filtered into four modulation-frequency ranges using a fourth-order, zero-phase Butterworth filter: 0.9–2.5 Hz, 2.5–5 Hz, 5–9 Hz and 12–40 Hz. These ranges were chosen to probe distinct temporal scales of speech information determined by prior computational modelling of the story content (Mandke et al., 2022): prosodic and phrase-level structure (0.9–2.5 Hz), syllabic timing (2.5–5 Hz), onset/segmental dynamics (5–9 Hz) and higher-frequency amplitude modulations (12–40 Hz). The resulting band-limited envelopes were stored and used as the speech regressors for all subsequent lagged coherence analyses.

### MRI acquisition and source modelling

Structural MRI scans were acquired on a Siemens Prisma 3T scanner using a T1-weighted MPRAGE sequence with 1-mm isotropic voxels. Anatomical scans were used for co-registration and individualised forward modelling, except where fsaverage was required. Cortical surface reconstructions were generated with FreeSurfer. For participants without individual MRIs, the fsaverage template was used, with the head shape-based co-registration to the MEG sensor space. A cortical source space was set up for each participant using an icosahedral subdivision (level 5), yielding an approximately uniform grid of dipoles on the cortical surface. Forward solutions were computed in MNE-Python using a hierarchical approach. When available, we used a subject-specific Boundary Element Method (BEM) solution. If only a BEM model was present, we generated the corresponding BEM solution. If no BEM files were available, we fitted a single-shell sphere model to the digitised headshape, slightly reducing the fitted radius (×0.98) to avoid intersection with the scalp. In all cases, we computed the forward solution using MEG sensors only with a minimum source-sensor distance of 5 mm. Noise covariance matrices were estimated from the first 120 s of the empty-room recording, and data covariance matrices from the first 600 s of the continuous story recording. We then constructed Linearly Constrained Minimum Variance (LCMV) beamformer filters using a regularisation parameter of 0.05 to select the orientation with maximum projected power at each source. These filters were applied to the continuous story data to obtain source time courses for each participant. To enable group-level analyses in a common cortical space, individual source estimates were morphed to fsaverage with 15 smoothing steps. For participants without individual MRI (analysed directly on fsaverage), the morphing step was omitted. All subsequent lagged coherence and ROI-level analyses were performed in fsaverage space.

### ROI definition and extraction

To summarise source-level signals within anatomically defined regions, we constructed bilateral ROIs on the fsaverage cortical surface by combining labels from the Desikan-Killiany parcellation (Desikan et al., 2006) and the Destrieux atlas (Destrieux et al., 2010). Using keyword-based matching of atlas label names, we defined ten bilateral ROIs targeting core nodes of the auditory and speech-language network: Heschl’s gyrus (HG), superior temporal gyrus (STG), middle temporal gyrus (MTG), superior temporal sulcus (STS), planum temporale (PT), inferior frontal gyrus pars opercularis (IFGop), inferior frontal gyrus pars triangularis (IFGtri), supramarginal gyrus (SMG), angular gyrus (AG) and inferior parietal lobule (IPL). For each ROI and hemisphere, we created a union label by merging the vertices of all atlas labels associated with that region.

For each participant, source-reconstructed time series were obtained at every cortical vertex in fsaverage space. ROI-level time series were derived by averaging the source-reconstructed signals across all vertices belonging to each union label. Vertex-wise metrics (including lagged coherence or connectivity measures) were reduced to the ROI level by averaging across all vertices within the corresponding label. This procedure yielded ROI-level time series and ROI-level summary measures for each hemisphere, participant and frequency band, which were used in subsequent lagged coherence and functional connectivity analyses.

### Lagged coherence computation

Speech-brain coupling was quantified using a lagged coherence metric computed at the source level for each modulation-frequency range. For each child and each band, the fsaverage-space source time series 𝑋 ∈ ℝ*^v^*^×*T*^ (𝑉 vertices × 𝑇 time points) and the corresponding band-limited speech envelope 𝑦 ∈ ℝ^#^ were first truncated to a common duration (about 10 minutes at 200 Hz). Both 𝑋 and 𝑦 were band-pass filtered using the same 4th-order zero-phase Butterworth filter applied during stimulus preprocessing. To characterise frequency-specific coupling, we obtained the analytic signal of the filtered sources and envelope using the Hilbert transform, yielding complex-valued time series 𝑍*_X_* and 𝑍*_y_*. For each source, we estimated complex coherency with the speech envelope as

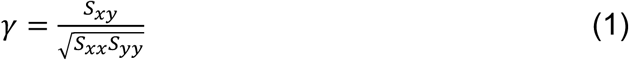

where 𝑆*_xy_*= ⟨𝑍*_X_* 𝑍^∗^*_y_*⟩, 𝑆*_xx_* = ⟨∣ 𝑍*_X_* ∣^+^⟩*_t_*, 𝑆*_yy_* = ⟨∣ 𝑍*_y_* ∣^+^⟩_*_ and ⟨⋅⟩_t_ denotes temporal averaging. We decomposed 𝛾 into real and imaginary parts (𝛾 = 𝑟𝑒 + 𝑖 𝑖𝑚) and defined lagged coherence as

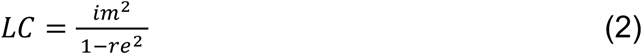

which selectively captures nonzero-lag coupling while suppressing instantaneous, zero-phase correlations that may arise from field spread or shared filtering. This yielded one lagged coherence value per source vertex and frequency band for each participant. The resulting maps were subsequently summarised within anatomically defined ROIs for statistical analysis.

### Statistical analysis of lagged coherence

Group differences in lagged coherence between typically-developing children and children with DLD were assessed at the ROI level. Lagged coherence was evaluated in four modulation-frequency bands (0.9–2.5 Hz, 2.5–5 Hz, 5–9 Hz and 12–40 Hz). For each frequency band and ROI, we compared lagged coherence values between groups using the Wilcoxon rank-sum test. Analyses were conducted separately for each ROI within each hemisphere. Statistical significance was defined as *p* < 0.05 (two-sided). In additional analyses, observed lagged coherence values were compared against permutation-derived chance-level distributions to verify that reported effects exceeded chance.

To estimate chance-level lagged coherence, we generated permutation-based null distributions using within-story circular time-shift surrogates. For each participant and frequency band, the band-limited speech envelope was circularly shifted by a random lag of at least 10 s, thereby disrupting temporal alignment. Lagged coherence was recomputed for each surrogate (500 permutations), averaged within ROIs, and then averaged across participants within each group to obtain group-level null distributions for each ROI and frequency band. Observed effects were considered reliable only when they exceeded the 95th percentile of the corresponding permutation-derived null distribution. Wilcoxon rank-sum results were interpreted only when observed group differences additionally exceeded the permutation-derived null model; isolated effects that did not exceed the null distribution were reported but not interpreted further.

### Functional connectivity computation using PLV

Functional connectivity between cortical regions was quantified using the PLV, computed at the source level between anatomically defined ROIs. Source time series were reconstructed using LCMV beamforming, morphed to fsaverage, and averaged within each ROI. PLV was then calculated between all ROI pairs for each participant and frequency band.

For each participant and frequency band, ROI time series were band-pass filtered using a zero-phase Butterworth filter and transformed to the analytic signal using the Hilbert transform. Instantaneous phase time series were extracted, and PLV between ROI i and ROI j was defined as:

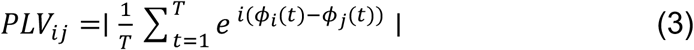

where 𝜙_t_(𝑡) and _j_(𝑡) denote the instantaneous phases of ROIs i and j, respectively, and 𝑇 is the number of time points. This yielded a symmetric PLV matrix for each participant and frequency band. PLV matrices were averaged across participants within each group (Controls and DLD) to obtain group-level connectivity estimates.

### Between-group permutation test for functional connectivity (Controls > DLD)

To assess group differences in connectivity, a non-parametric, one-sided permutation test was applied independently to each ROI pair and frequency band. The observed test statistic was the difference in group-mean PLV (averaged across participants within each group):

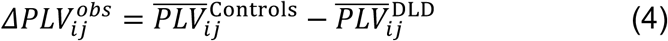

Group labels were then randomly permuted at the participant level while preserving group sizes, and the PLV difference was recomputed for each permutation. This procedure was repeated 𝑁_CD>E_ = 5000 times, generating a null distribution of PLV differences expected under the hypothesis of no group effect. For each ROI pair (i, j), the one-sided p-value was computed as:

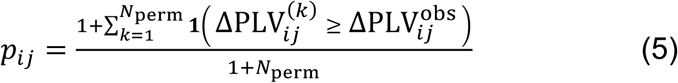

where 𝟏(⋅) is an indicator function. Edges with *p* < 0.05 were considered statistically significant, indicating stronger phase synchronisation in Controls relative to the DLD group. No correction for multiple comparisons was applied, consistent with an exploratory network-level analysis.

### Within-group permutation test (PLV > chance)

To determine whether observed PLV values exceeded those expected by chance within each group, a second non-parametric permutation test was performed using circular time-shift surrogates. For each participant, the phase time series of each ROI was circularly shifted by a random amount (independently for each ROI), thereby destroying inter-regional phase relationships while preserving the temporal autocorrelation and spectral properties of the signals. PLV matrices were recomputed for each surrogate dataset and averaged across participants within each group. This procedure was repeated 𝑁_CD>E_ = 1000 times, yielding a null distribution of group-mean PLV values under the null hypothesis of no true inter-regional coupling. For each ROI pair, a one-sided p-value was computed as the proportion of surrogate PLV values exceeding the observed group-mean PLV.

## Acknowledgements

The authors would like to thank all the participants involved in the study. This research was funded by a donation to UG from the Yidan Prize Foundation. The sponsor played no role in the study design, data interpretation nor writing of the report.

## Author contributions

MK: Conceptualisation, Data Acquisition, Data Curation, Methodology, Data Analysis, Visualisation, Writing – original draft. GF: Data Acquisition, Writing – Review and editing. SR: Data Acquisition, Writing – Review and editing. LP: Data Acquisition, Writing – Review and editing. UG: Conceptualisation, Project Administration, Funding Acquisition, Resources, Supervision, Writing – original draft.

## Data and code availability

All preprocessing, source reconstruction and lagged coherence computations were implemented in Python using MNE-Python and custom scripts, with MATLAB used for statistical analyses. De-identified data and analysis code are available from the corresponding author upon reasonable request, subject to institutional and ethical constraints.

